# Combinatory therapy targeting mitochondrial oxidative phosphorylation improves efficacy of IDH mutant inhibitors in acute myeloid leukemia

**DOI:** 10.1101/749580

**Authors:** Lucille Stuani, Marie Sabatier, Feng Wang, Nathalie Poupin, Claudie Bosc, Estelle Saland, Florence Castelli, Lara Gales, Camille Montersino, Emeline Boet, Evgenia Turtoi, Tony Kaoma, Thomas Farge, Nicolas Broin, Clément Larrue, Natalia Baran, Marc Conti, Sylvain Loric, Pierre-Luc Mouchel, Mathilde Gotanègre, Cédric Cassan, Laurent Fernando, Guillaume Cognet, Aliki Zavoriti, Mohsen Hosseini, Héléna Boutzen, Kiyomi Morita, Andrew Futreal, Emeline Chu-Van, Laurent Le Cam, Martin Carroll, Mary A. Selak, Norbert Vey, Claire Calmettes, Arnaud Pigneux, Audrey Bidet, Rémy Castellano, Francois Fenaille, Andrei Turtoi, Guillaume Cazals, Pierre Bories, Yves Gibon, Brandon Nicolay, Sébastien Ronseaux, Joe Marszalek, Courtney D. DiNardo, Marina Konopleva, Yves Collette, Laetitia K. Linares, Floriant Bellvert, Fabien Jourdan, Koichi Takahashi, Christian Récher, Jean-Charles Portais, Jean-Emmanuel Sarry

## Abstract

Isocitrate dehydrogenases (IDH) are involved in redox control and central metabolism. Mutations in IDH induce epigenetic and transcriptional reprogramming, differentiation bias, BCL-2 dependence and susceptibility to mitochondrial inhibitors in cancer cells. Here we show that high sensitivity to mitochondrial oxidative phosphorylation (OxPHOS) inhibitors is due to an enhanced mitochondrial oxidative metabolism in cell lines, PDX and patients with acute myeloid leukemia (AML) harboring IDH mutation. Along with an increase in TCA cycle intermediates, this AML-specific metabolic behavior mechanistically occurs through the increase in methylation-driven CEBPα- and CPT1a-induced fatty acid oxidation, electron transport chain complex I activity and mitochondrial respiration in IDH1 mutant AML. Furthermore, an IDH mutant inhibitor that significantly and systematically reduces 2-HG oncometabolite transiently reverses mitochondrial FAO and OxPHOS gene signature and activities in patients who responded to the treatment and achieved the remission. However, at relapse or in patients who did not respond, IDH mutant inhibitor failed to block these mitochondrial properties. Accordingly, OxPHOS inhibitors such as IACS-010759 improve anti-AML efficacy of IDH mutant inhibitors alone and in combination with chemotherapy *in vivo*. This work provides a scientific rationale for combinatory mitochondrial-targeted therapies to treat IDH mutant-positive AML patients, especially those unresponsive to or relapsing from IDH mutant-specific inhibitors.

---

Changes in intermediary and energy metabolism provide the flexibility for cancer cells to adapt their metabolism to meet energetic and biosynthetic requirements for proliferation^1–4^. Manipulating glycolysis, glutaminolysis, fatty acid β-oxidation (FAO) or oxidative phosphorylation (OxPHOS) markedly reduces cell growth *in vitro* and *in vivo* and sensitizes acute myeloid leukemia (AML) cells to chemotherapeutic drugs^5–13^. The importance of the metabolic reprogramming in this disease is further illustrated by recurrent mutations in genes of two crucial metabolic enzymes, isocitrate dehydrogenases (IDH) 1 and 2, present in more than 15% of AML patients^14–17^.

The impact of IDH mutation and the related accumulation of the oncometabolite (R)-2-hydroxyglutarate (2-HG) have been well documented in leukemic transformation and AML biology^18–28^. As IDH mutations are early events in oncogenesis and are systematically conserved at relapse^29–31^, IDH1/2 mutated (IDHm) enzymes represent attractive therapeutic targets and small molecules specifically inhibiting the mutated forms of these enzymes have been developed and recently approved by the FDA^32–41^. Both the IDH2m- and IDH1m-inhibitors promote differentiation and reduce methylation levels as well as significantly decrease 2-HG levels^33,36,37,42,43^. Overall response rates for ivosidenib (IDH1m inhibitor) and enasidenib (IDH2m inhibitor) are highly encouraging up to 30 or 40% in monotherapy in phase I/II clinical trials for newly diagnosed or relapsed/refractory AML patients respectively). However, several mechanisms of resistance to these targeted therapies have been already identified^37–39,42,44^. Moreover, suppression of serum 2-HG level alone did not predict response in patients, as many non-responders also displayed a significant decrease in the amount of 2-HG^37,40,42,44–47^. Importantly, multiple pathways involved in signaling, clonal heterogeneity or second-site mutation are very recently considered to be responsible for relapses in patients treated with IDH mutant inhibitors^38,48–50^. Thus, targeting IDH mutant activity is not sufficient to achieve a durable clinical response in most patients and new combinatory approaches need to be designed.

Given the crucial roles of wild type (WT) IDH1/2 in cell metabolism (e.g. Krebs cycle, OxPHOS, cytosolic and mitochondrial redox, anabolism including lipid biosynthesis) and in human disease^51^, a better understanding of the contribution of oncogenic IDH mutations to metabolism and metabolic homeostasis is expected to lead to new therapeutic strategies. Several studies have demonstrated that IDH mutant cancer cells exhibit some metabolic specificities^52–58^. However, none of these studies have definitively shown how metabolic changes elicited by IDH mutations modulate cell proliferation and drug resistance or impact therapeutic response in AML. In particular, the role of metabolism in resistance to IDHm inhibitors has not been yet comprehensively studied in AML. Although existing literature in the field described several vulnerabilities to mitochondrial inhibitors in IDH1/2-mutant cells from solid tumors and AML^9,59–62^, no studies have also fully demonstrated why IDH mutant cells are more sensitive to mitochondrial inhibitors in AML. We therefore hypothesized that mitochondrial oxidative phosphorylation plays a crucial role in IDH mutant biology and in the response of AML patients with IDH mutation to IDHm inhibitors.

In the present study, we perform multi-omics and functional approaches using 2 engineered AML cell lines, 12 PDX models from two clinical sites (Toulouse Hospital and University of Pennsylvania) and 123 patient samples from four clinical sites (Toulouse Hospital TUH, Bordeaux Hospital BUH, Marseille Hospital IPC and MD Anderson Cancer Center MDACC) to test this hypothesis and to expressly understand the mitochondrial reprogramming induced by IDH1 mutation and its role in the response to IDH mutant specific inhibitors.

## Results

### A higher susceptibility of IDH1 mutant AML cells to mitochondrial inhibitors is due to their enhanced OxPHOS activity

First we confirmed a higher sensitivity of IDH1/2-mutant cells from primary AML patient samples (WT, n=64; MUT, n=56; TUH, BUH, IPC, MDACC; Supplementary Table 1) and two genetically diverse cell lines to mitochondrial inhibitors such as OxPHOS inhibitors, including a new electron transport chain (ETC) complex I inhibitor IACS-010759^63^ and metformin, ETC complex III (antimycin A, AA; atovaquone, ATQ), ETC complex V (oligomycin, OLIGO) or BCL2 inhibitors (ABT-199, ABT-263) (Fig. 1a-c and Supplementary Fig. 1a). Interestingly, their 2-HG response was heterogeneous (Fig. 1d). Whereas metformin induced an increase in 2-HG content, the BCL2 inhibitor ABT-199 caused a reduction in the amount of the oncometabolite. This strongly suggests an enhancement of mitochondrial metabolic dependency in IDH mutant subgroup of AML patients without a systematic correlation with 2-HG content.

**Figure 1.**
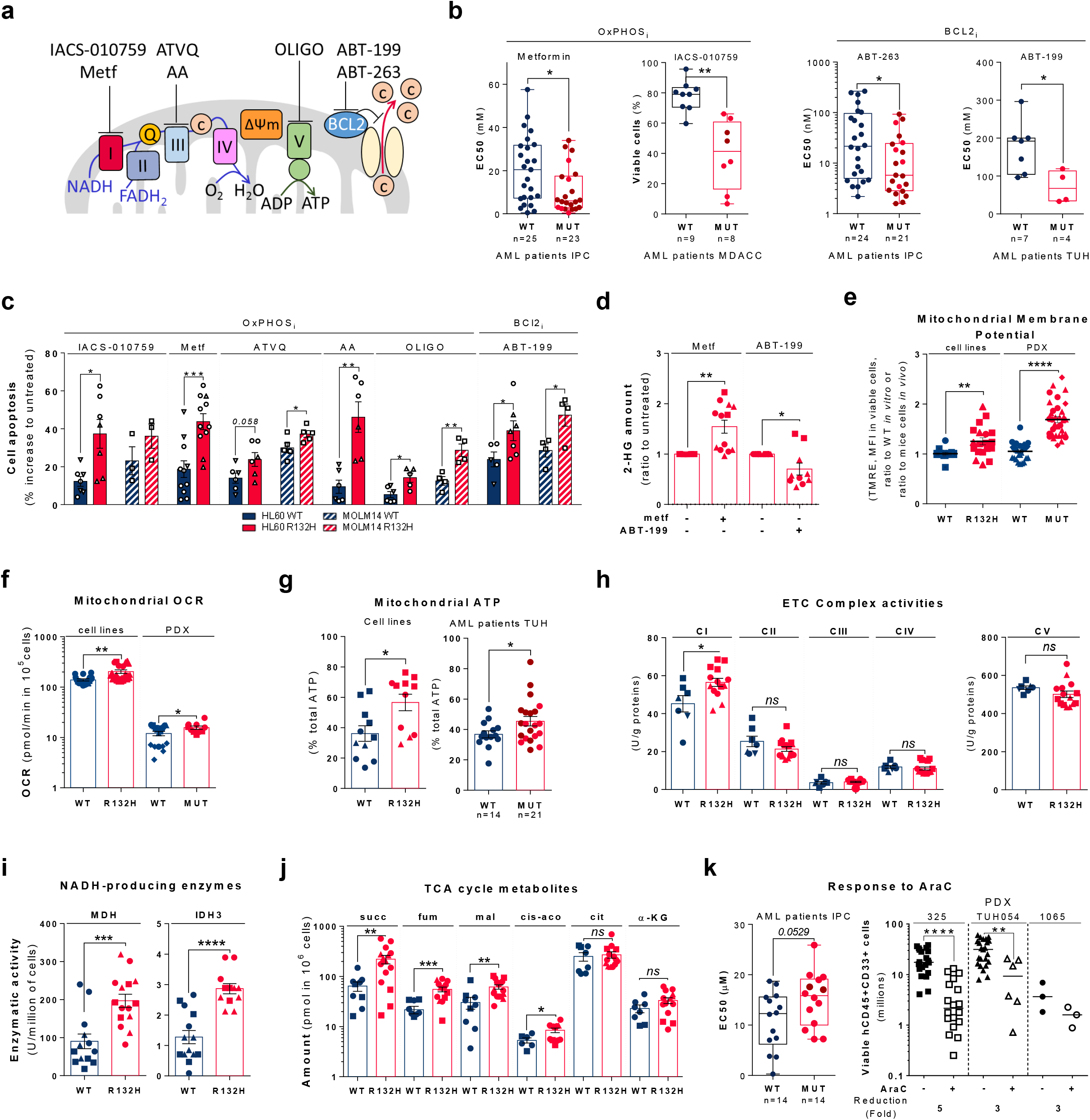
IDH1 mutant cells exhibit a higher susceptibility to OxPHOS_i_ and BCL2_i_ is linked to their enhanced mitochondrial capabilities and OxPHOS activity in AML. **(a)** Schematic representation of the electron transport chain (ETC) and BCL2 with OXPHOSi and BCL2i used in this study (Metf= metformin; AA=antimycin A; ATVQ=atovaquone; oligo=oligomycin). **(b)** Plots of EC50 values from ATP viability assays of metformin and ABT-263 after 48h, from annexin V positive cells assays of ABT-199 after 24h and percent of viable cells after 72h of IACS-010759 in primary samples with WT or MUT IDH1 (red circles) or IDH2 (burgundy circles). See also Supplementary Table 1 for patient information. **(c)** Apoptosis induction following IACS-010759 (100nM during 48h for HL60 and during 6 days for MOLM14), 48h metformin (10mM), antimicyn A (10μM), atovaquone (20μM for HL60 and 40 μM for MOLM14), oligomycin (2μM) and ABT-199 (200nM) in HL60 and MOLM14 IDH1 WT or R132H. Errors bars indicate Mean±SEM of at least three independent experiments. **(d)** 2-HG fold change following 24h treatment with metformin (10mM) and ABT-199 (200nM) in HL60 and MOLM14 IDH1 WT or R132H. Errors bars indicate Mean±SEM of at least two independent experiments. **(e)** TMRE mitochondrial potential assay in HL60 and MOLM14 IDH1 WT or R132H measured *in vitro* (n≥3) and *in vivo* in PDX (3 patients IDH1 WT and 3 patients IDH1 MUT) to estimate Mitochondrial Membrane Potential (MMP). See also Supplementary Table 1 for patient information. Errors bars indicate Mean±SEM. **(f)** Mitochondrial Oxygen Consumption Rate (OCR) of HL60 and MOLM14 IDH1 WT or R132H measured *in vitro* (n≥3) and *ex vivo* in PDX after cell-sorting (4 patients IDH1 WT and 2 patients IDH1 MUT). See also Supplementary Table 1 for patient information. Errors bars indicate Mean±SEM. **(g)** Mitochondrial ATP in HL60 and MOLM14 IDH1 WT and R132H (n≥3) and in patients with IDH WT (n=14) or MUT IDH1 (red circles) or IDH2 (burgundy circles) (n=21). See also Supplementary Table 1 for patient information. Errors bars indicate Mean±SEM. **(h)** Mitochondrial ETC complex activities in HL60 and MOLM14 IDH1 WT and R132H. Errors bars indicate Mean±SEM of at least three independent experiments. **(i)** NADH-producing enzyme activities of malate dehydrogenase (MDH) and isocitrate dehydrogenase (IDH3) in HL60 and MOLM14 IDH1 WT and R132H. Errors bars indicate Mean±SEM of at least three independent experiments. **(j)** Succinate (succ), fumarate (fum), malate (mal), cis-aconitate (cis-aco), citrate (cit) and α-KG amounts measured over 24h culture in HL60 and MOLM14 IDH1 WT and R132H. Errors bars indicate Mean±SEM of at least three independent experiments. **(k)** Plots of EC50 values of AraC determined from ATP viability assays at 48h (left panel) and total number of human viable AML cells expressing CD45 and CD33 in AraC-treated compared with PBS-treated IDH1 mutant AML-xenografted mice in bone marrow and spleen (right panel). See also Supplementary Table 1 for patient information. For each panel **(b–k)**, HL60 IDH1 WT are represented in blue by circles (clone 4), triangles up (clone 2) and triangles down (clone 7) whereas R132H are represented in red by circles (clone 11) and triangles up (clone 5). MOLM14 are represented by squares, blue for IDH1 WT and red for IDH1 R132H (both induced by doxycycline). Groups were compared with unpaired two-tailed t test with Welch’s correction. *p<0.05; **p<0.01; ***p<0.001; ns, not significant.

To better understand why IDH1 mutant cells have a higher sensitivity to mitochondrial inhibition, we extensively analyzed several biochemical, enzymatic and functional features relative to mitochondrial activity in IDH1 mutant *versus* WT AML cells from two genetically diverse AML cell lines *in vitro* and six patient-derived xenografts *in vivo* (Supplementary Fig. 1b). Mitochondrial membrane potential, oxygen consumption, ATP-linked respiration and ATP content were all significantly enhanced in IDH mutant AML cells *in vitro* and *in vivo* (Fig. 1e-g and Supplementary Fig. 1c-e). Importantly, ETCI complex (and not other ETC complexes) activity, NADH-producing enzyme activity of TCA enzymes such as malate dehydrogenase (MDH2) and isocitrate dehydrogenase (IDH3) and concentration of Krebs cycle intermediates (except α-KG) were also increased in IDH1 mutant AML cells (Fig. 1h-j), indicating an increase in mitochondrial NADH availability, mitochondrial activities and OxPHOS dependency specifically in IDH1 mutant AML cells. Interestingly, this was not due to an increase in mitochondrial biogenesis as shown by mitochondrial mass, protein content of ETC complexes, citrate synthase activity or ratio between mitochondrial and nucleic DNA, which were not affected (Supplementary Fig. 1f-j).

Drug-resistant cancer cells have recently shown to be enriched in cells exhibiting a high OxPHOS signature and enhanced mitochondrial function in several cancers including myeloid malignancies^11,12,64,65^. Accordingly, we observed that IDH1 mutant cells were more resistant to conventional cytarabine (AraC) chemotherapy than IDH1 WT cells *in vitro* and in three PDX models *in vivo* that are low responders (Fig. 1k and Supplementary Fig. 1k), as previously defined to distinguish patients with high from low AraC response *in vivo*^11^.

### Methylation- and CEBPα- dependent mitochondrial FAO is increased in IDH1 mutant cells

In order to further identify the mitochondrial reprogramming induced by IDH1 mutation, we next performed a computational analysis of the metabolic network of IDH1 mutant cells based on human genome scale metabolic network reconstruction Recon2 (7440 metabolic reactions)^66^. To reconstruct active leukemic metabolic networks of IDH1 WT and mutant AML cells at a global level, we integrated transcriptomic data and applied metabolic constraints according to metabolite production and consumption rates measured in the corresponding cell culture supernatants (exometabolome)^67,68^ (Fig. 2a). This analysis identified a significant enrichment of active reactions in various carbon metabolic pathways (N-glycan synthesis, fructose and mannose metabolism, dicarboxylate metabolism; Fig. 2a) in IDH1 mutant cells and predicted a major change in FAO in these cells (Fig. 2a-b), especially CPT1 which is required to initiate the transfer of fatty acids from the cytosol to the mitochondrial matrix for oxidation (Supplementary Fig. 2a). This prediction prompted us to assess key features of FA utilization in AML cells and patients with IDH1 mutation. First, we measured acyl-CoAs as readout of FA catabolism. As expected, acetyl-CoA, succinyl-CoA, free coenzyme A and FA oxidation rate were significantly increased in IDH1 mutant AML cells (Fig. 2c-d).

**Figure 2.**
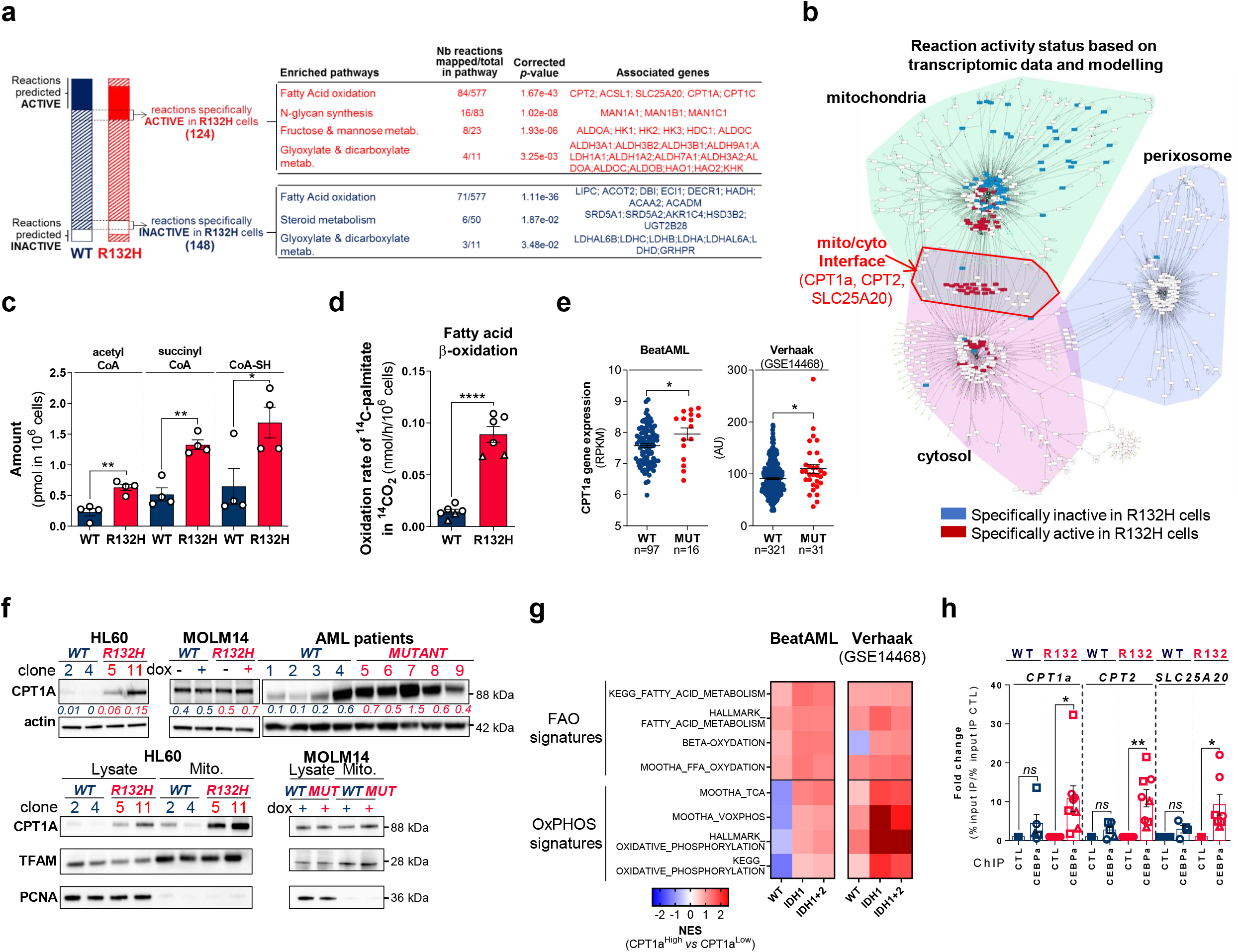
Methylation- and CEBPα- dependent mitochondrial fatty acid oxidation is increased in IDH1 mutant cells. **(a)** Comparison of the predicted activity of reactions in the metabolic network of HL60 IDH1 WT *vs*. R132H cells. Predictions of reactions activity or inactivity was made using the Recon 2 metabolic network reconstruction and transcriptomic data from HL60 IDH1 WT and R123H^23^. Pathway enrichment was performed on the set of reactions identified as specifically active (red) or specifically inactive (blue) in R123H cells. Corrected *p*-values were obtained by performing a hypergeometric test followed by a Bonferroni correction. **(b)** Visualization of modulated reactions within the fatty acid oxidation pathway of the Recon 2 metabolic network. Reactions predicted to be specifically active (red) or inactive (blue) in R123H cells based on transcriptomic data alone (left panel) or using the computational modeling approach (right panel) were mapped using the MetExplore webserver^94,95^. **(c)** Acetyl-CoA, succinyl-CoA, free coenzyme A (CoASH) amounts measured over 24h culture in HL60 IDH1 WT clone 4 and R132H clone 11 lysates. Errors bars indicate Mean±SEM of two independent experiments with 2 technical replicates for each analyzed with unpaired two-tailed t test with Welch’s correction. **(d)** ^14^C palmitate oxidation by HL60 IDH1 WT clone 4 (circle) and 2 (triangle) and R132H clone 11 (circle) and 5 (triangle) to assess β-oxidation rate. Errors bars indicate Mean±SEM of six independent experiments analyzed with unpaired two-tailed t test with Welch’s correction. **(e)** *CPT1a* gene expression across AML patient samples from GSE14468 (Verhaak cohort) and BeatAML^96^ datasets in function of their IDH1 status. Groups were compared using unpaired non-parametric Mann-Whitney test. **(f)** Total lysates (left panel) and lysates of purified mitochondria (mito) (right panel) of HL60 and MOLM14 IDH1 WT and R132H and total lysates of primary samples IDH1 WT or MUT (bottom left panel) were immunoblotted with the indicated antibodies. See also Supplementary table 1 for patient information. **(g)** Normalized enrichment score (NES) following GSEA analysis of patients with high or low expression of CPT1a (mediane as the reference) in IDH WT, IDH1 mutant or IDH1+2 mutant across AML transcriptomes from two-independent cohorts, BeatAML and Verhaak (GSE14468). **(h)** qChIP experiments showing the relative recruitment of CEBPα on *CPT1a, CPT2* and *SLC25A20* locus in mutant IDH1 R132H *versus* IDH1 WT HL60 and MOLM14, as indicated. Results were represented as the relative ratio between the mean value of immunoprecipitated chromatin (calculated as a percentage of the input) with the indicated antibodies and the one obtained with a control irrelevant antibody. HL60 IDH1 WT are represented in blue by circles (clone 4) and triangles (clone 2) whereas R132H are represented in red by circles (clone 11) and triangles (clone 5). MOLM14 are represented by squares, blue for IDH1 WT and red for IDH1 R132H (both induced by doxycycline). Errors bars indicate Mean±SEM of at least two independent experiments analyzed with unpaired two-tailed t test with Welch’s correction. *p<0.05; **p<0.01; ***p<0.001; ****p<0.0001; ns, not significant.

The regulation of FAO is complex and involves different signaling pathways and allosteric regulation (Supplementary Fig. 2a). We first examined AMP kinase (AMPK), a master regulator of energy and metabolic homeostasis. Surprisingly, whereas the AMPK protein level was increased in IDH1 mutant cell lines and patients, its activation by phosphorylation was not increased (Fig. 2b). Furthermore, we did not observe an increase in the AMP/ATP ratio in mutants compared to WT (Supplementary Fig. 2c), suggesting that AMP, the primary allosteric activator of AMPK, does not directly activate AMPK in these cells. No increase in ADP/ATP ratio was detected, suggesting that neither AMP nor ADP enhanced the canonical phosphorylation of AMPK Thr172 *via* LKB1 in IDH1 mutant AML cells (Supplementary Fig. 2c). These data showed that AMPK is not activated directly by AMP or *via* phosphorylation in mutant cells, indicating that the changes seen in FAO in mutant cells reflect an AMPK-independent mechanism. Similarly, the PKA pathway did not appear to be differentially activated and to be involved into the FAO regulation (Supplementary Fig. 2d). Interestingly, protein levels and phosphorylation of both acetyl-CoA carboxylases (ACC1 and ACC2) that regulate malonyl-CoA level and hence modulate mitochondrial FA shuttling and oxidation, were decreased in IDH1 mutant AML cells and therefore favor FAO (Supplementary Fig. 2e). Transcriptomic data using two independent AML cohorts (GSE 14468^69^ and TCGA^70^) reinforced this observation demonstrating a significant decrease in *ACACA* (coding ACC1) and *ACACB* (coding ACC2) mRNA expression in IDH1 mutant cells (Supplementary Fig. 2f). Moreover, previous reports have shown that gene expression is closely modulated by histone and DNA methylation in IDH mutant cells^18,71,72^. To address the importance of histone methylation at the *ACACA* promoter, we performed quantitative chromatin immunoprecipitation (qChIP) experiments to measure the levels of trimethylation of lysine 4 of histone H4 (H3K4me3) and of trimethylation of lysine 27 of histone H3 (H3K27me3), two epigenetic markers associated with transcriptional activation and repression, respectively. While we did not observe any differences in H3K4me3 occupancy of the *ACACA* promoter, a significant increase in H3K27me3 on *ACACA* occurred specifically in IDH1 mutant cells (Supplementary Fig. 2g). Furthermore, we performed a gene set enrichment analysis (GSEA) with a curated FAO gene signature^73^ and found that this signature was enriched in IDH1 mutant AML cells (Supplementary Fig. 2h). The most highly expressed gene in IDH1 mutant cells was the key component of FA shuttling into mitochondria *CPT1a*. Consistent with this, *CPT1a* and its isoform *CPT1b* mRNA levels were also significantly upregulated in our IDH1 mutant cell lines and in two independent AML cohorts (Fig. 2e and Supplementary Fig. 2i). Also, CPT1A protein was significantly increased in total cell lysates and in mitochondria isolated from IDH1 mutant cells compared to IDH1 WT cells and in IDH1 mutant primary samples (Fig. 2f). Finally, GSEA analysis comparing the transcriptomes of AML patients with IDH WT, IDH1 or IDH1/2 mutation revealed higher enrichment of FA metabolism and OxPHOS gene signatures in CPT1a^HIGH^ patients with IDH mutations in two-independent cohorts (Fig. 2g). This strengthens the observation that CPT1a plays a crucial role in FA metabolism and OxPHOS in IDH mutant AML cells.

We previously demonstrated that IDH1 mutation and its 2-HG product dysregulate CEBPα^23^, a well-known transcriptional regulator of several genes involved in glucose and lipid metabolism. Moreover, CEBPα was the second most highly expressed gene of the FAO gene signature in IDH1 mutant cells (Supplementary Fig. 2h). Thus, we performed qChIP assays to assess CEBPα binding to promoters of genes encoding FA transporters. We observed that the recruitment of endogenous CEBPα to promoter of *CPT1a, CPT2*, and *SLC25A20* that mediates the transport of acyl-carnitines of different length across the mitochondrial inner membrane from the cytosol to the mitochondrial matrix, increased specifically in IDH1 mutant cells (Fig. 2h). Furthermore, CEBPα silencing led to a reduction of mitochondrial basal OCR as well as ATP-linked and FAO-coupled OCR in IDH1 WT and to a greater extent in IDH1 mutant AML cells (Supplementary Fig. 2j-k). Together, these results indicate that IDH1 mutant cells display a gene signature specific for FA shuttling and a high FAO activity in CPT1a- and CEBPα- dependent manner to support mitochondrial activity.

### Reduction of 2-HG with IDHm inhibitors transiently reverses the mitochondrial phenotype of IDH mutant AML cells

As IDH mutant cells exhibited higher mitochondrial activity than WT cells, we investigated the impact of IDHm inhibitors (FDA approved ivosidenib AG-120 and its preclinical version AGI-5198 for IDH1 and enasidenib AG-221 for IDH2) on this OxPHOS phenotype. Because Phase I clinical trials NCT02074839 or NCT01915498 showed 40% overall response rate for ivosidenib or enasidenib monotherapy for IDH mutant AML patients with relapsed or refractory AML, respectively^38,39^, we reasoned that non-responders in this study might have different mitochondrial and FAO status. Interestingly, we performed comparative transcriptomic analyses of IDH mutant patients characterized as good responders to IDHm inhibitor at complete remission (bone marrow blast < 5% and normalization to peripheral blood count; n=6 patients) *versus* before treatment and at relapse post-IDHm inhibitor *versus* at complete remission. We thus shown that curated gene signature related to OxPHOS, Krebs cycle, FAO and pyruvate metabolism were enriched in patients before treatment and upon relapse (Fig. 3a). Furthermore, these gene signatures were significantly enriched after IDHm inhibitor treatment in two IDH mutant patients who did not respond to IDHm inhibitor (Fig. 3a). In particular, the expression of the FAO genes *CPT1a, CPT2, SLC25A20* was significantly reduced in good responders to IDHm inhibitor at complete remission compared to before treatment but then increased at relapse to reach the same level as before treatment (Fig. 3b). In two AML cell lines harboring IDH1 mutation, AG-120 and AGI-5198 treatments significantly reduced 2-HG levels and decreased the expressions of *CEBPα, CPT1a, CPT2* and *SLC25A20* (Supplementary Fig. 3a-b). Furthermore, IDH1m inhibitors prevented the recruitment of endogenous CEBPα to promoter of *CPT1a, CPT2* and *SLC25A20* (Supplementary Fig. 3c). As FAO is one of the major biochemical pathways that support OxPHOS and mitochondrial function especially in AML^5,11,74^, it was surprising to observe that IDH1m inhibitor maintained or even increased FAO-coupled (Fig. 3c), basal mitochondrial (Fig. 3d) and ATP-linked OCR (Fig. 3e). We then assessed several mitochondrial activities after treatment with IDHm inhibitors. IDH1m inhibitors also maintained or increased mitochondrial response in both IDH1 mutant cell lines such as their mitochondrial membrane potential (Supplementary Fig. 3d), TCA cycle intermediate concentrations (Fig. 3f), ETC complex activities or protein amounts (Supplementary Fig. 3e-g). Altogether, these results demonstrated that, while decreasing the level of 2-HG and some FAO features, IDHm inhibitors only transiently reverse OxPHOS phenotype, leading us to specifically consider combination with OxPHOS inhibitor for these patients.

**Figure 3.**
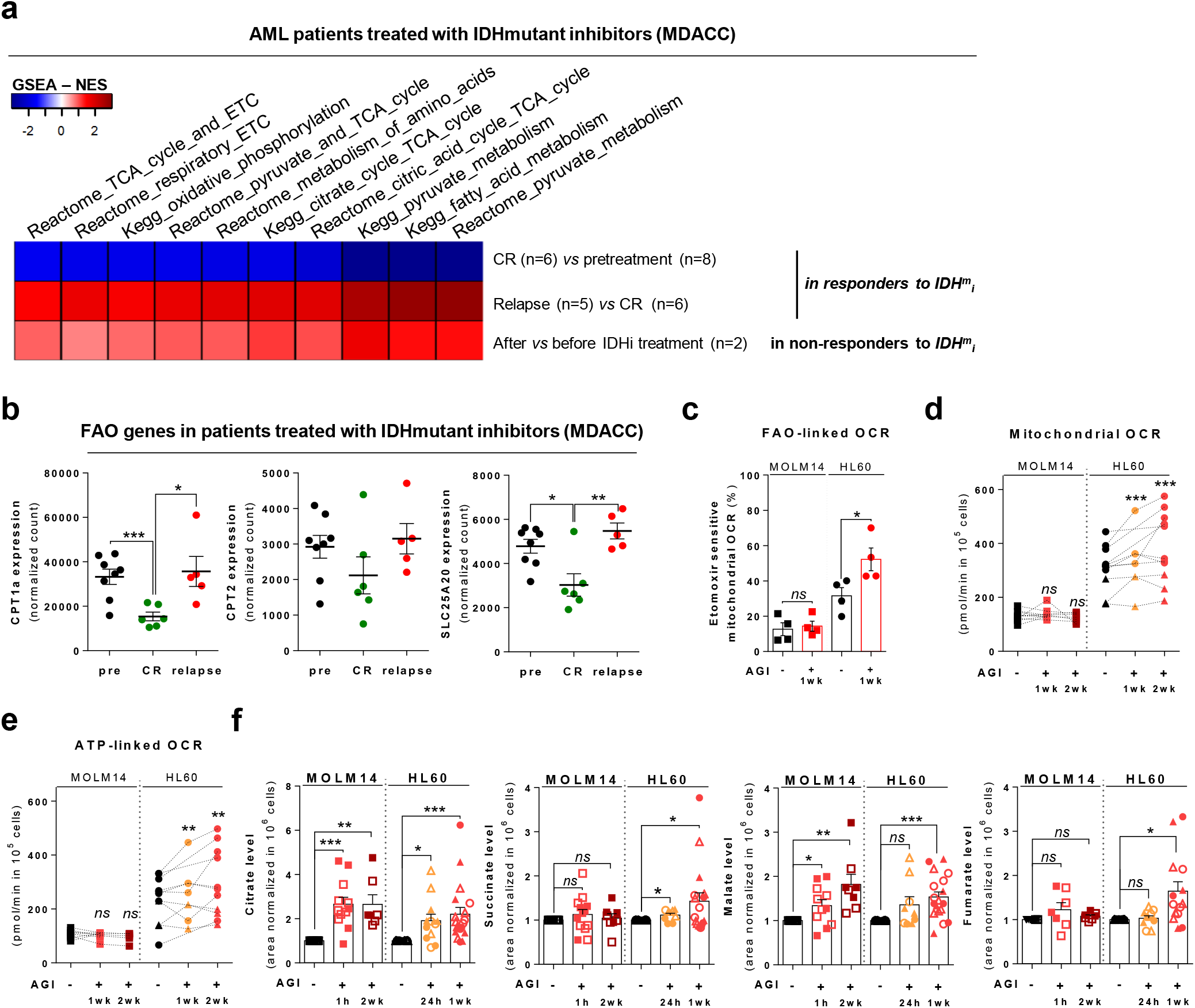
IDH mutant inhibitors reverse 2-HG production but do not necessarily decrease high OxPHOS phenotype and mitochondrial metabolism. **(a)** GSEA Normalized Enrichment Score (NES) from several transcriptomic signatures of AML patients with IDH mutation characterized as responders to IDHm inhibitor before treatment (8 patients), at complete remission CR (6 patients) and at relapse (5 patients); or as non-responders to IDHm inhibitor (2 patients) before and after treatment with IDHm inhibitor. See also Supplementary table 1 for patient information. **(b)** Relative mRNA levels for *CPT1a, CPT2*, and *SLC25A20* in AML patients with IDH mutation characterized as responders to IDHm inhibitor before treatment (8 patients), at complete remission CR (6 patients) and at relapse (5 patients). **(c)** FAO-linked OCR in HL60 and MOLM14 IDH1 R132H following 1-week treatment with AGI-5198 (2μM). Errors bars indicate Mean±SEM of four independent experiments. **(d)** Mitochondrial OCR of HL60 and MOLM14 IDH1 R132H in vehicle (DMF) and after 24h, 1 week or 2 weeks treatment with AGI-5198 (2μM). Errors bars indicate Mean±SEM of at least three independent experiments. **(e)** ATP-linked OCR of HL60 and MOLM14 IDH1 R132H in vehicle (DMF) and after 24h, 1 week or 2 weeks treatment with AGI-5198 (2μM). Errors bars indicate Mean±SEM of at least three independent experiments. **(f)** Citrate, succinate, malate and fumarate levels normalized to internal standard measured over 24h culture in HL60 and MOLM14 IDH1 R132H following 24h, 1 week or 2 weeks treatment with AGI-5198 (2μM-plain symbols) or AG-120 (1μM-empty symbols). HL60 IDH1 R132H are represented in red by circles (clone 11) and triangles (clone 5), whereas MOLM14 IDH1 R132H are represented by red squares. Errors bars indicate Mean±SEM of at least three independent experiments. For panels **(c–g)**, groups were compared with unpaired two-tailed t test with Welch’s correction. *p<0.05; **p<0.01; ***p<0.001; ns, not significant.

### Treatment with OxPHOS inhibitors enhances anti-leukemic effects of IDH-mutant specific inhibitors alone and in combination with cytarabine

We assessed several mono, duplet or triplet therapeutic approaches *in vivo* with IDHm inhibitor AG-120 (150 mg/kg, twice every day, 3 weeks), ETC complex I inhibitor IACS-010759 (8 mg/kg, every other day, 3 weeks) or/and AraC (30 mg/kg, every day, 1 week) in two IDH1 R132 PDX models with variable engraftment capacity (high engrafting patient #325 in Fig. 4a-b, Supplementary Fig. 4; low engrafting patient #1065 in Supplementary Fig. 5a). The level of 2-HG was greatly reduced in the PDX sera upon all therapies compared to control group while α-KG remained unchanged in PDX #325 (Fig 4c and Supplementary Fig. 4a). Total cell tumor burden was significantly reduced in mono, duplet and triplet therapy with a greater effect in the triplet therapy cohort compared to vehicle cohort or AraC monotherapy (Fig. 4d). Similarly, apoptosis was also increased in all treatments except AG-120 mono and duplet therapy with IACS and we observed a greater significance in the triplet therapy compared to vehicle (Supplementary Fig. 4b). Expression of the myeloid differentiation marker CD15 was also significantly increased in duplet therapy combining AraC and IACS and even more in the triplet therapy (Fig 4e and Supplementary Fig. 4c). Interestingly, mitochondrial OxPHOS function assessed *in vivo* by mitochondrial membrane potential was only decreased in the triplet therapy (Supplementary Fig. 4d). Furthermore, mitochondrial ATP content and respiratory capacities were increased after AraC or AG-120 alone or combined together but rescued with the addition of IACS in the triplet therapy *in vivo* (Fig. 4f and Supplementary Fig. 4e-g). Analysis of mice serum metabolomes showed that aspartate level was significantly reduced in all AraC groups, in particular in the duplet therapy with AG-120 and the triplet therapy. Lactate level was enhanced in all groups with IACS including the triplet therapy as key biomarker of IACS-010759 response (Fig. 4g). Similar *in vivo* experiments with lower engrafting IDH1-R132 PDX showed a lower level in anti-leukemic and biological effects of the mono, duplet and triplet combinations (Supplementary Fig. 5). However and more importantly in this low responder PDX, the triplet therapy induces a greater decrease in the total cell tumor burden, in mitochondrial activity through decreased mitochondrial membrane potential, mitochondrial ATP and enhanced lactate amount in mice sera (Supplementary Fig. 5b-e). Of note, global toxicity of the triplet therapy or duplet therapies with AraC was primarily driven by AraC toxicity (Supplementary Fig. 4h-i and Fig. 5f-g). Altogether, these results not only confirmed that IDHm inhibitor does not necessarily reverse metabolic and mitochondrial (especially, enhanced OxPHOS phenotype) features of IDH mutant cells *in vivo* but also that combining this drug with ETC complex I inhibitor in presence or not of standard AraC chemotherapy increases its drug efficacy *in vivo*, notably by inducing a Pasteur effect (e.g. increased lactate in response to the inhibition of mitochondrial ATP production). Finally, taking advantage of this preclinical study, we have identified a set of metabolic changes (here called AML metabolic profiling) that represent a combination of classic hallmarks of the Pasteur effect and other metabolic adaptations to predict the response to IDH *plus* OxPHOS inhibitors (Supplementary Fig. 6 and Supplementary Table 2) and to monitor the efficacy of their response (Supplementary Table 3) in IDH mutant AML subgroup.

**Figure 4.**
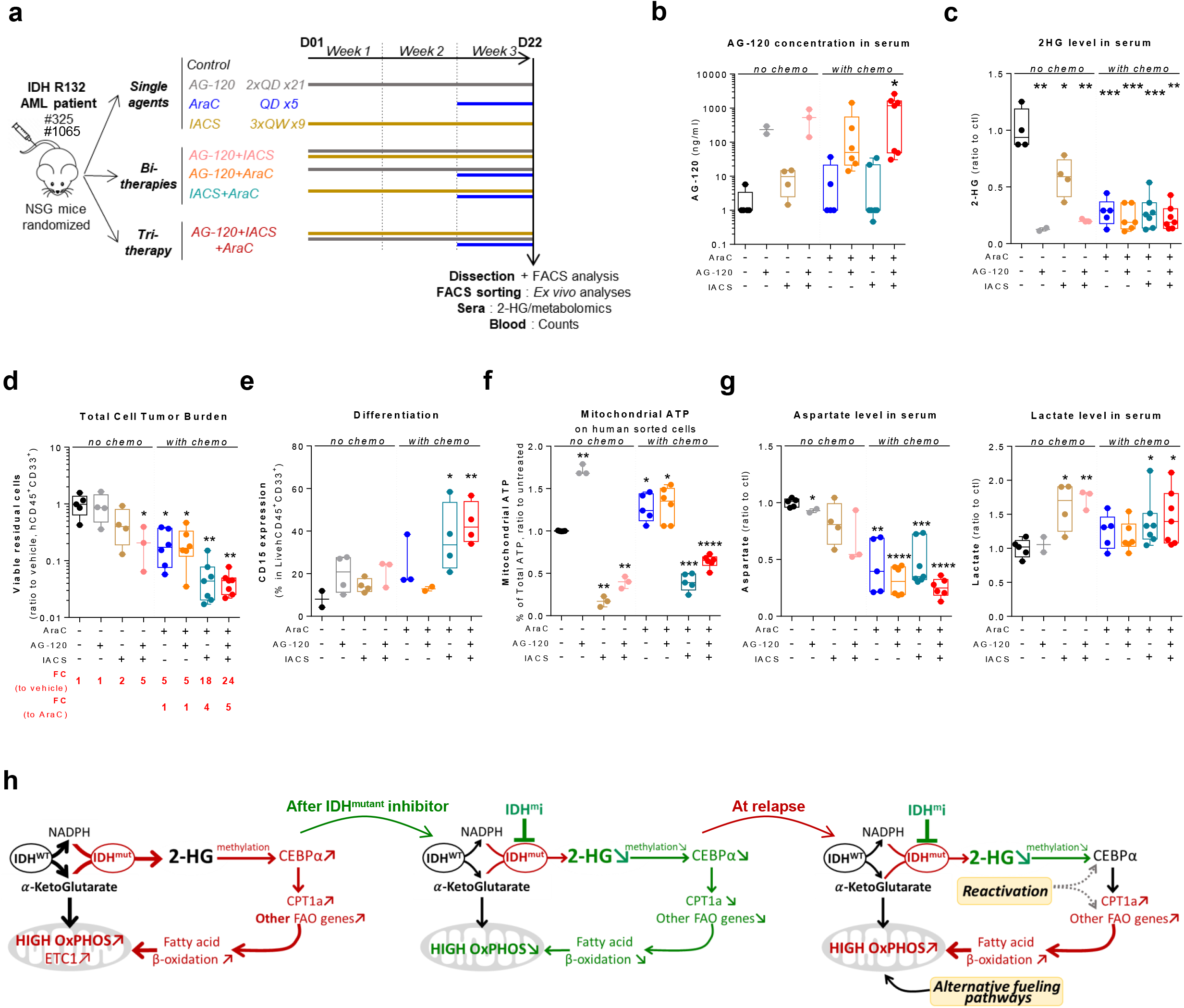
Treatment with an OxPHOS inhibitor enhances anti-leukemic effects of IDH-mutant specific inhibitors alone and in combination with cytarabine in a high engrafter IDH1 mutant AML PDX model. **(a)** Experimental scheme detailing administration time of intraperitoneal AraC, IACS 010759 and AG-120 by gavages in PDX. See also Supplementary table 1 for patient information. **(b)** AG-120 concentration in mice sera of PDX 325. **(c)** 2-HG level normalized to control group in sera of IDH1 R132 PDX 325 mice after mono-, duplet- or triplet-therapies compared with vehicle. **(d)** Total number of human viable AML cells expressing CD45 and CD33 in mono-, duplet- or triplet-therapies compared with vehicle and AraC-treated IDH1 R132 PDX 325 mice in bone marrow and spleen. Fold change (FC) between each group and the mean of vehicle or AraC group. **(e)** Percent of human viable cells expressing CD15 in bone marrow in mono-, duplet- or triplet-therapies compared with vehicle-treated IDH1 R132 PDX 325 mice. **(f)** Percent of mitochondrial ATP contribution to total ATP after FACS-sorting of human viable cells expressing CD45 and CD33 in bone marrow of IDH1 R132 PDX 325 mice treated with mono-, duplet- or triplet-therapies compared with vehicle. **(g)** Aspartate and lactate levels normalized to control group in mice sera of IDH1 R132 PDX 325. **(h)** Schematic diagram of metabolic reprogramming induced by IDH1 mutation in AML cells and its impact on OxPHOS status through FAO regulation in absence of treatment, after treatment with IDHm inhibitor then at relapse. For panels **(b–g)**, groups were compared with unpaired two-tailed t test with Welch’s correction. *p<0.05; **p<0.01; ***p<0.001; ****p<0.0001; ns, not significant.

## Discussion

Since the discovery of IDH mutations in various cancers, significant efforts have been directed to understand the extent to which these oncogenic mutations directly impact metabolism, histone/DNA methylation and gene expression^18,26,27,75–78^, cell proliferation and differentiation bias^19,21–23,34^. However, questions remain unanswered related to the impact of IDH mutation on mitochondrial energetic metabolism in IDH mutant cells. Here we address several aspects of these different crucial points in IDH mutant cell biology and explain mitochondrial dependency in the basal condition and upon IDHm inhibitor treatment. IDH mutation induces mitochondrial reprogramming that contributes to maintain the pool of α-KG to support 2-HG production and to replenish other Krebs cycle intermediates necessary for anabolic reactions, oxygen consumption and ATP production by oxidative phosphorylation in AML (Fig. 4h). Importantly, treatment with IDHm inhibitors transiently reverses FAO and OxPHOS activities that are maintained or enhanced in non-responders or at relapse (Fig. 4h).

Several studies in different cancers have consistently shown increased mitochondrial phenotypes in IDH mutant cells^59,61,79,80^. Our study helps to explain the dependence of IDH mutant cells on mitochondria and energetic metabolism required to sustain mutant cell proliferation, in particular for synthesis of α-KG and NADPH, the substrates of mutant IDH enzyme activity. It is noteworthy that disturbances in cellular and mitochondrial metabolism can contribute to the chemoresistance of AML cells^5,6,11^. Here we showed that IDH1 mutant AML cells exhibited a higher OxPHOS phenotype than WT cells, consistent with a lower response to AraC. We also observed a OxPHOS hyperactivity of IDH mutant cells and confirmed a strong ETC1 complex dependency in AML^7,11,63,81,82^. Accordingly, mutant AML cells exhibit enhanced vulnerabilities to various small molecules targeting mitochondrial OxPHOS such as inhibitors of ETC complex I, III and V. Consistent with literature reports and current clinical trials, we also observed that IDHm AML cells were more sensitive to BCL2 inhibition by ABT-199^20,62,83^. Several combinations of BCL2i with newly approved targeted therapies such as FLT3i and IDHi are under clinical assessment (NCT03735875 and NCT03471260, respectively). Interestingly, 2-HG levels did not correlate with apoptosis in these experiments, as its concentration decreased after ABT-199 while it increased after metformin treatment, a result already observed in IDH1 R132H transformed mammary epithelial cells^61^. This lack of correlation between cellular concentration of 2-HG, cell survival and sensitivity to various inhibitors is of particular interest as it has been shown that neither did inhibitors of mutant IDH reverse all IDH-mutant phenotypes nor did suppression of 2-HG alone predict response to IDHm inhibitors^33,44,45,59,84,85^. Our data and previous reports clearly show that 2-HG is essential but insufficient in mediating and mimicking all the complex metabolic consequences of IDH mutation^56,86^. These observations strongly suggest that innovative combinatory therapies might be useful in this patient subgroup. Of particular interest, whereas IDHm inhibitors maintain or increase mitochondrial and OxPHOS activity in IDH mutant cell lines and primary patients *in vitro, in vivo* and in relapsed or refractory AML patients in clinical trials, triplet combination using IDH1m inhibitor, OxPHOS inhibitor (such as IACS-010759), and AraC showed promising effects in IDH1 mutant AML PDX. Therefore, this triplet drug combination may represent a beneficial alternative for AML patients unresponsive to IDH mutant-specific inhibitors.

In this regard, transcriptomic analysis before administration of an IDHm inhibitor in AML patients with IDH mutation, at complete remission and at relapse and in patients who did not respond to the drug revealed an enrichment in FAO and OxPHOS gene signatures at relapse and in non-responders. In particular, expression of several genes participating in FAO such as *CPT1a, CPT2* and *SLC25A20* correlate with relapse and response to IDHm inhibitor and could potentially be used in clinics to predict and monitor the therapeutic efficacy of IDHm and OxPHOS inhibitors. Moreover, we observed that lactate concentration in mice sera was not modified and that mitochondrial ATP was maintained or increased by treatment with IDH1m inhibitor alone while duplet or triplet therapies with OxPHOSi lead to significant increase in lactate concentration and reduction in mitochondrial ATP in our high responder PDX harboring IDH1 mutation. Accordingly, we proposed an AML metabolic profiling based on measuring mitochondrial ATP, OxPHOS and FAO gene signatures from primary cells, and lactate, aspartate and 2-HG in the sera of patients. This could represent a potentially powerful tool in identifying and understanding dependence on individual mitochondrial FAO and OxPHOS activities. Consequently, this combination of metabolic and genetic approaches can also be used to predict and monitor responses to the duplet or triplet therapy combining IDHi and OxPHOSi (Supplementary Fig. 6, Supplementary Table 2 and Supplementary Table 3) in the context of the functional precision cancer medicine^87,88^.

Of particular interest, it was very recently shown that for patients with IDH1 or IDH2 mutation who responded to IDHm inhibitors in clinics and then relapsed, acquired resistance to this molecularly targeted therapy was caused by the emergence of clones with a second-side IDH mutation in the wild type IDH allele without the initial IDH mutation, rescuing 2-HG production^48^. This reinforced the therapeutic interest/potential of our combinatory strategy. Finally, our study supports the merit of future clinical trials testing the combination of IDHm inhibitors and mitochondrial inhibitors with cytarabine treatment. Because this proposed therapeutic strategy will overcome different newly identified mechanisms of resistance to IDH mutant inhibitors^38,48,49^, this would be especially relevant as alternative therapeutic approaches for the treatment of those patients that are not unresponsive to or relapsing from IDH mutant-specific inhibitors.

## Methods

### Primary AML samples

Primary AML patient specimens are from five clinical sites [University of Pennsylvania (UPENN), Philadelphia, PA; MD Anderson Cancer Center at University of Texas, Houston, (MDACC), Toulouse University Hospital (TUH), Toulouse, France, Institut Paoli-Calmettes (IPC), Marseille, France and Bordeaux University Hospital (BUH), Bordeaux, France].

<u>For TUH and BUH</u>, frozen samples of bone marrow or peripheral blood were obtained from patients diagnosed with AML after signed informed consent in accordance with the Declaration of Helsinki, and stored at the HIMIP collection (BB-0033-00060) and CRB-K BBS (BB-0033-00036). According to the French law, HIMIP biobank collection and BBS biobank have been declared to the Ministry of Higher Education and Research (DC 2008-307, collection 1 for HIMIP and DC 2014-2164 for BBS) and obtained a transfer agreement (AC 2008-129) after approbation by the Comité de Protection des Personnes Sud-Ouest et Outremer II (ethical committee). Clinical and biological annotations of the samples have been declared to the CNIL (Comité National Informatique et Libertés ie Data processing and Liberties National Committee). <u>For IPC clinical site</u>, *ex vivo* drug sensitivity was performed on previously frozen (HEMATO-BIO-IPC 2013--015 clinical trial, NCT02320656) or fresh (CEGAL-IPC-2014-012, NCT02619071 clinical trial) mononuclear cell samples from 49 AML patients after informed consent (Supplementary Table 1). Both trials have been approved by a properly constituted Institutional Review Board (Comité de Protection des Personnes) and by the French National Security Agency of Medicine and Health Products (ANSM). The samples are subjected to NGS to screen for mutations within a selected panel of ~150 actionable genes in AML (i.e.) known to be of prognostic value and/or druggable. <u>For UPENN</u>, AML samples were obtained from patients diagnosed with AML in accordance with U.S. Common Rules at the Stem Cell and Xenograft Core Facility at the UPENN School of Medicine and with informed consent in accordance with institutional guidelines. Peripheral blood or bone marrow samples were frozen in FCS with 10% DMSO and stored in liquid nitrogen. The percentage of blasts was determined by flow cytometry and morphologic characteristics before purification. <u>For MDACC cohort</u>, bone marrow samples were collected from the 11 AML patients who were treated with AG-120 (n=6) and AG-221 (n=5) in The University of Texas MD Anderson Cancer Center from clinical trials NCT01915498 (enasidenib AG-221 for IDH2 mutated patients), NCT02074839 (ivosidenib AG-120 for IDH1 mutated patients). All patients had provided written informed consent for sample collection and subsequent data analysis. Nine patients had achieved complete remission (responders; n = 9), and 2 patients never achieved remission (non-responders; n=2). For a subset of responders, longitudinal samples were obtained at pre-treatment (n = 8), complete remission (n = 6), and at the time of relapse (n = 5). For non-responders (n=2), we analyzed the paired bone marrow samples obtained at pre-treatment and post-treatment. These samples were analyzed by targeted capture next generation sequencing using SureSelect custom panel of 295 genes (Agilent Technologies, Santa Clara, CA, USA), as well as RNA sequencing. Bone marrow morphology and karyotyping data were interpreted by the board certified hematopathologists.

### Mice and mouse xenograft model

Animals were used in accordance with a protocol reviewed and approved by the Institutional Animal Care and Use Committee of Région Midi-Pyrénées (France). NOD/LtSz-SCID/IL-2Rγchainnull (NSG) mice were produced at the Genotoul Anexplo platform at Toulouse (France) using breeders obtained from Charles River Laboratories. Mice were housed in sterile conditions using high-efficiency particulate arrestance filtered microisolators and fed with irradiated food and sterile water.

Human primary AML cells were transplanted as reported previously^11,89–91^. Briefly, mice (6–9 weeks old) were sublethally treated with busulfan (20 mg/kg/day) 24 hours before injection of leukemic cells. Leukemia samples were thawed at room temperature, washed twice in PBS, and suspended in Hank’s Balanced Salt Solution at a final concentration of 0.2-10×10^6^ cells per 200μL of Hank’s Balanced Salt Solution per mouse for tail vein injection. Transplanted mice were treated with antibiotic (Baytril) for the duration of the experiment. Daily monitoring of mice for symptoms of disease (ruffled coat, hunched back, weakness, and reduced mobility) determined the time of killing for injected animals with signs of distress. If no signs of distress were seen, mice were initially analyzed for engraftment 8 weeks after injection except where otherwise noted.

### *In vivo* mice treatment

Eight to 18 weeks (PDX) after AML cell transplantation and when mice were engrafted (tested by flow cytometry on peripheral blood or bone marrow aspirates), NSG mice were treated as described below:

- <u>AraC treatment</u>: NSG mice were treated by daily intraperitoneal injection of 60 mg/kg AraC for 5 days; AraC was kindly provided by the pharmacy of the TUH. For control, NSG mice were treated daily with intraperitoneal injection of vehicle, PBS 1X.
- <u>IACS-10759 treatment</u>: IACS-10759 was solubilized in water containing 0.5% methylcellulose before administration to mice. NSG mice were treated 3 times a week by gavage of 8 or 5 mg/kg IACS-10759 (according to weight loss of mice) for 21 days. For control, NSG mice were treated by daily gavage of vehicle. IACS-10759 was kindly provided by Dr. Joe Marszalek.
- <u>AG-120 treatment</u>: AG-120 (Ivosidenib, AGIOS Pharmaceuticals) was solubilized in water containing 0.5% methylcellulose and 0.2% Tween80 before administration to mice. NSG mice were treated twice a day by gavage of 150mg/kg AG-120 for 21 days. For control, NSG mice were treated twice a day by gavage of vehicle.

Mice were monitored for toxicity and provided nutritional supplements as needed.

### Assessment of Leukemic Engraftment

Assessment of leukemic engraftment was measured as reported previously^11^. Briefly, NSG mice were humanely killed in accordance with European ethics protocols. Bone marrow (mixed from tibias and femurs) and spleen were dissected in a sterile environment and flushed in Hank’s Balanced Salt Solution with 1% FBS. MNCs from peripheral blood, bone marrow, and spleen were labeled with hCD33-PE (555450), mCD45.1-PerCP-Cy5.5 (156058), hCD45-APC (5555485), and hCD44-PECy (7560533) (all antibodies from BD Biosciences) to determine the fraction of human blasts (hCD45^+^mCD45.1^-^hCD33^+^hCD44^+^ cells) using flow cytometry. All antibodies used for cytometry were used at concentrations between 1/50 and 1/200 depending on specificity and cell density. Analyses were performed on a CytoFLEX flow cytometer with CytoExpert software (Beckman Coulter) and FlowJo 10.2 (Tree Star). The number of AML cells/μl peripheral blood and number of AML cells in total cell tumor burden (in bone marrow and spleen) were determined by using CountBright beads (Invitrogen) using described protocol.

### Statistical analyses

Statistical analyses were conducted using Prism software v6.0 (GraphPad Software, La Jolla, CA, USA). For *in vitro* and *in vivo* studies, statistical significance was determined by the two-tailed unpaired Student’s t-test. For transcriptomic analysis of cohorts, statistical significance was determined by the non-parametric Mann-Withney test. A pvalue < 0.05 was considered statistically significant. For all figures, ns, not significant, *p%0.05, **p%0.01, ***p%0.001, ****p%0.0001. Unless otherwise indicated, all data represent the mean ± standard error of the mean (SEM) from at least three independent experiments. Box-and-whisker plots displays all the individual data points as well as the corresponding median. For metabolomic analysis, Seahorse and ATP assays, each biological replicates represents the mean of at least two technical replicates.

## Data availability statement

RNAseq data from Fig. 3 are part of a clinical trial and available upon request.

## Acknowledgements

We thank The Cancéropoles PACA and GSO, the Network MetaboCancerGSO, Karine Marendziak, all members of mice core facilities (UMS006, ANEXPLO, Inserm, Toulouse) for their support and technical assistance, and Audrey Sarry, Prof. Véronique De Mas and Eric Delabesse for the management of the Biobank BRC-HIMIP (Biological Resources Centres-Inserm Midi-Pyrénées “Cytothèque des hémopathies malignes”) that is supported by CAPTOR (Cancer Pharmacology of Toulouse-Oncopole and Région). We thank Anne-Marie Benot, Muriel Serthelon and Stéphanie Nevouet for their daily help about the administrative and financial management of the Sarry lab. We thank the Institut Paoli Calmettes Direction de la Recherche Clinique et de l’Innovation (DRCI), the Hematology Department, and the IPC/CRCM Biological Resource Center for sample collection and processing. We also thank patients and their families.

This work was also supported by grants from the Région Midi-Pyrénées (FLEXAML, J-E.S.), the Plan Cancer Biologie des Systèmes 2014 (FLEXAML; J-E.S.), the Laboratoire d’Excellence Toulouse Cancer (TOUCAN; contract ANR11-LABEX), the Canceropole GSO (MetaboCancerGSO; J-E.S., J-C.P., L.L.C.) the Programme Hospitalo-Universitaire en Cancérologie (CAPTOR; contract ANR11-PHUC0001), La Ligue Nationale de Lutte Contre le Cancer, the Fondation ARC, the Fondation Toulouse Cancer Santé and the Association G.A.E.L. MetaTOUL (Metabolomics & Fluxomics Facitilies, Toulouse, France, www.metatoul.fr) and LEMM are part of the national infrastructure MetaboHUB-ANR-11-INBS-0010 (The French National infrastructure for metabolomics and fluxomics, www.metabohub.fr). MetaToul is supported by grants from the Région Midi-Pyrénées, the European Regional Development Fund, the SICOVAL, the Infrastructures en Biologie Santé et Agronomie (IBiSa, France), the Centre National de la Recherche Scientifique (CNRS) and the Institut National de la Recherche Agronomique (INRA). The project has been partly supported by Canceroprole PACA, SIRIC (grant INCa-Inserm-DGOS 6038 2012-2017), Canceropôle-SIRIC EmA, INCA 2017-024-COLLETTE and INCa 2017-024-MAL. C. M. was supported by SIRIC (grant INCa-Inserm-DGOS 6038 2012-2017) and Canceropôle-SIRIC EmA. E. T. was supported by SIRIC-Montpellier Cancer Program (grant INCa-Inserm-DGOS 6045 2012-2017). N. Ba. and M. K. were supported by funding from Leukemia Spore grant P50 (CA100632-16).

## Author contributions

L.S., M.S. and J-E.S designed experiments. L.S., M.S., C.B., E.S., T.F., N.Br., C.L., N. Ba., M. Co., S.L., C.Cas., G.Co., A.Z., M.H., H.B., L.K.L. performed *in vitro* experiments. M.S., C.B., E.S., E.B., T.F., P-L.M, M.G. performed *in vivo* experiments. C.M. and R.C. performed chemogrammes’ analysis. L.S., F.C., L.G., E.T., E.C-V, A.T. and G. Ca. performed metabolomics analyses. N.P., L.F. and F.J. performed genome-scale metabolic network analysis. L.S., T.F. and T.K. performed transcriptomic analysis on publically available cohorts. F.W., K.M., A.F. and K.T. performed and analyzed RNAseq analysis on patient samples treated with IDHm inhibitors. AML bone marrow and blood samples were provided by M.Ca. (UPENN), N.V. and Y.C. (IPC), C.Cal., A.P. and A.B. (BUH), P.B. and C.R. (TUH), C.D.D and M.K. (MDACC). J. M. and M.K provided IACS-010759. B.N. and S.R. provided and measured AG-120 in mice sera. L.L.C., F.F., A.T., Y.G., J.M., C.D.D, M.K., L.K.L, F.B. and J-C.P managed the resources and shared their expertise. L.S. and J-E.S. wrote the manuscript and designed the figures. N.P., M.A.S., F.F., F.J., C.R. and J-C.P reviewed the manuscript. J-E-S directed the research.

## Competing interests statement

C.D.D. is a consultant for Agios and Celgene, and served on the advisory board for Bayer, Karyopharm, MedImmune, and AbbVie. S.R. and B.N. are both employees and own stock in Agios Pharmaceuticals.

The rest of the authors declare no competing interests.

